# MicroFinder: Conserved gene-set mapping and assembly ordering for manual curation of bird microchromosomes

**DOI:** 10.1101/2025.05.09.653066

**Authors:** Thomas C. Mathers, Michael Paulini, Cibele G. Sotero-Caio, Jonathan M. D. Wood

**Affiliations:** Tree of Life, Wellcome Sanger InsAtute, Wellcome Genome Campus, Hinxton, Cambridge, CB10 1SA, UK

## Abstract

Obtaining chromosomally complete genome assemblies across the tree of life is a major goal of biodiversity genomics. However, some lineages remain recalcitrant to assembly despite recent advances in sequencing technologies and assembly tools. Birds present a substantial genome assembly challenge due to the presence of tiny, hard to assemble microchromosomes that are often highly fragmented or even missing in draft genome assemblies. As such, bird genomes require a large amount of expert manual curation effort via manipulation of genome-wide Hi-C contact maps and many current chromosome-level bird genome assemblies do not resolve the known karyotype. Microchromosomes have distinct genetic and epigenetic features. They are GC-biased, gene-rich, highly methylated, and have distinct spatial organisation in the centre of the nucleus. Importantly, they are conserved across avian evolution. Here, using a reference set of expert curated bird genomes, we have identified a set of conserved microchromosome genes and developed MicroFinder, a pipeline that uses this gene set to find small microchromosome fragments in draft genome assemblies to act as anchors for manual curation of microchromosomes. We demonstrate how MicroFinder can be used to improve the speed and accuracy of bird genome curation. Furthermore, we highlight the usefulness of MicroFinder by carrying out MicroFinder-enabled re-curation of 12 previously released chromosome-scale bird genome assemblies, increasing the sequence content of microchromosome models.

## Introduction

Recent advances in sequencing technology have dramatically improved the quantity, quality and taxonomic breadth of reference genome assemblies across the tree of life (Feron & Waterhouse, 2022; Lewin et al., 2001; Rhie et al., 2021; The Darwin Tree of Life Project Consortium, 2021). Automated assembly of accurate long reads followed by scaffolding with high throughput *in vivo* chromatin conformation capture sequence data (Hi-C) and manual curation (Howe et al., 2021) routinely results in genome assemblies that meet or exceed accepted gold standard metrics (Lawniczak et al., 2022). However, some lineages are recalcitrant to assembly and challenges remain to generate complete, chromosomally resolved genome assemblies for all taxa (H. Li & Durbin, 2024).

Within vertebrates, birds present a substantial assembly challenge due to the presence of tiny, hard to assemble, microchromosomes. Since early cytogenetic studies, it has been recognised that bird genomes typically contain six to eight pairs of large macrochromosomes and 31 to 33 pairs of small microchromosomes (Tegelström & Ryttman, 1981). In chicken, macrochromosome size based on a near-T2T assembly ranges from 250 Mb to 30 Mb, and microchromosomes range from 23 Mb to 2.5 Mb (Huang et al., 2023). Ten of the smallest microchromosomes (ranging in size from 6.8 to 2.5 Mb) are further categorised as “dot” chromosomes based on their minute size, morphology and extensive pericentromeric heterochromatin. Once considered unimportant DNA fragments (Newcomer, 1955, 1957), cytogenetics and genomics have revealed that microchromosomes are highly conserved across avian evolution and contain many important and highly expressed housekeeping genes (Liu et al., 2021; van Brink, 1959; Waters et al., 2021). Furthermore, microchromosomes have distinct genetic and epigenetic features setting them apart from macrochromosomes: they are GC-biased, gene-rich, highly methylated, and have distinct spatial organisation in the centre of the nucleus (Habermann et al., 2001; McQueen et al., 1998; O’Connor et al., 2019; Perry et al., 2021; Smith et al., 2000).

Most recent bird genome assembly projects follow the Vertebrate Genome Project (VGP) assembly pipeline which uses accurate PacBio HiFi long reads for *de novo* assembly combined with Hi-C data for long range scaffolding and phasing (Larivière et al., 2024). This pipeline produces assemblies with excellent contiguity and completeness statistics. However, these metrics do not fully capture the challenge of assembling the smallest bird chromosomes as they represent a small fraction of the total sequence content. Strikingly, despite high-quality sequence data, bird genome assemblies often do not fully resolve the known karyotype (**Figure 1a**; **Supplementary Table 1**). Of 105 species with chromosome-scale genome assemblies in International Nucleotide Sequence Database Collaboration (INSDC) databases that also have karyotype data, 62 (59 %) differ from the expected karyotype by 2 or more chromosomes, with the majority (57/62) having fewer chromosomes than expected. Primarily, this is due to failure to assemble and identify the full set of microchromosomes (Barros et al., 2023; Peona, Blom, et al., 2021) and even in karyotype-resolved assemblies, microchromosomes are often highly fragmented and can be incomplete (M. Li et al., 2022). Painstaking manual curation of bird genomes after *de novo* assembly and scaffolding is therefore an essential assembly step. For example, the Hi-C contact map for the draft genome assembly of the pink-footed goose *Anser brachyrhynchus* (assembled by the Darwin Tree of Life (DToL) project [Lopez Colom & O’Brien, 2024]) reveals 28 clear chromosomal elements (**Figure 1b**), yet closely related karyotyped geese all have 40 or 41 chromosomes (Uno et al., 2019; Wójcik & Smalec, 2007).

**Figure 1.**
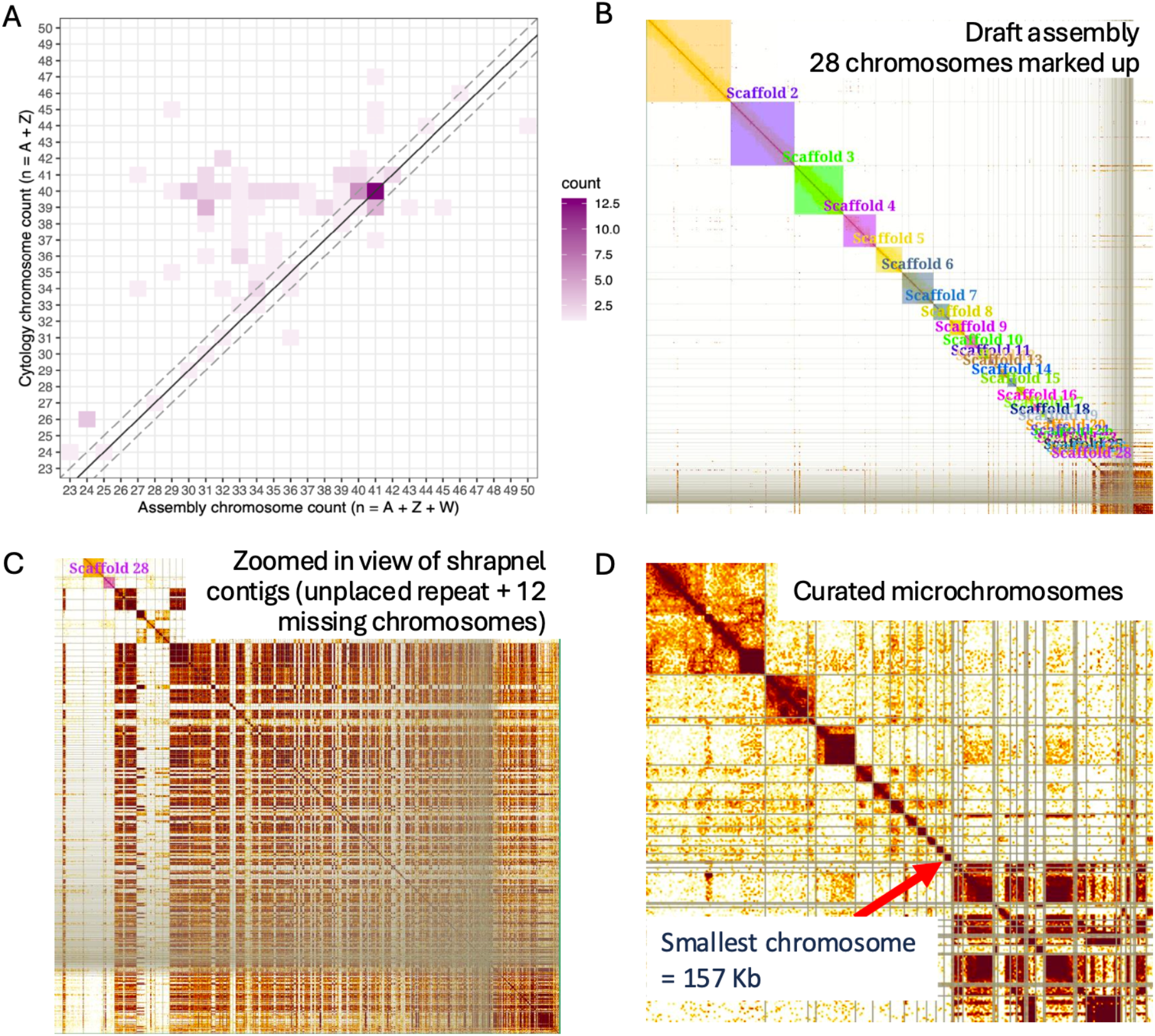
Bird genome assemblies are often not karyotype-complete and require extensive manual curation. (**A**) Correspondence analysis of chromosome counts in chromosome-scale genome assemblies versus their respective haploid karyotype for 105 bird species. Colour gradient reflects the number of species in each category (bin of assembly (x) versus karyotype (y) count). The Solid black line marks the match of the chromosome number in assemblies (y-axis) and predicted chromosome number using cytology (x-axis). The dashed diagonal lines indicate ± 1 chromosome margin of error to account for expected variation from assemblies of males (homogametic sex will usually have 1 less assembled chromosome). (**B**) Hi-C contact map for the draft genome assembly of *Anser brachyrhynchus* (assembled by the Darwin Tree of Life (DToL) project [Lopez Colom & O’Brien, 2024]). Coloured squares highlight 28 clear chromosomal elements identified during an initial assembly curation (painted “Scaffolds” in PretextView). (**C**) A zoomed in view of the unplaced assembly content grouped at the bottom-right of image (**B**). (**D**) Hi-C contact map of the curated *A. brachyrhynchus* genome assembly zoomed in on the smallest 11 chromosomes. Content to the right of the red arrow is unplaced content. Microchromosomes have elevated background Hi-C signal but appear as independent elements in the Hi-C map.

Therefore, at least 12 chromosomes are expected to be among the unplaced “shrapnel” content located at the bottom right of the Hi-C contact map which predominantly contains repetitive sequence (**Figure 1C**). To resolve the assembly, genome curators sift through shrapnel scaffolds to identify and assemble microchromosome fragments (**Figure 1d**). Techniques include making use of the elevated Hi-C background signal between microchromosomes (due to their central position in the nucleus), genome alignments with reference species and mapping of protein coding genes. This process is slow and laborious and there is a high likelihood of sequence content being missed from the assembled chromosomes.

Here, to aid manual curation of bird genomes, we took advantage of conserved gene content to identify microchromosome fragments in draft genome assemblies. Using 11 high-quality, manually curated bird genomes generated as part of the VGP, 25 Genomes Project and DToL (The Darwin Tree of Life Project Consortium, 2021), as well as a near telomere-to-telomere (T2T) assembly of chicken (Huang et al., 2023), we identified a set of conserved microchromosome genes and have developed MicroFinder (https://github.com/sanger-tol/MicroFinder), a pipeline that uses this gene set to find candidate microchromosome contigs from draft assemblies to improve the speed and accuracy of manual curation. Using this approach, we revisited 12 previously released bird genome assemblies and improved the content and representation of their assembled microchromosomes.

## Results and Discussion

### Identification of conserved microchromosome genes

Given the gene-dense nature of microchromosomes and their conserved synteny across birds, we hypothesised that a dense marker set of protein coding genes would enable the identification of microchromosome fragments in draft genome assemblies. To generate a set of marker genes, we made use of expert-curated genome assemblies generated for the VGP, DToL and 25 Genomes projects. We selected 11 published genome assemblies with NCBI RefSeq or Ensemble rapid release gene-sets (**Supplementary Table 2**). We also included a recent, near-T2T assembly of chicken (Huang et al., 2023). Together, these 12 assemblies span nine bird orders and 11 families (**Supplementary Table 2, Figure 2A**). Of note, this collection includes three high confidence genome assemblies (bCucCan1, bTaeGut1 and GGswu, herein referred to as the *ToL reference set*) that are commonly used by genome curators at the Welcome Sanger Tree of Life (ToL) program as references for whole genome alignments when curating new bird assemblies. Additionally, six of the selected assemblies have been confirmed to be karyotype-complete based on cytology (**Supplementary Table 2, Figure 2A**). Of the remaining assemblies, two species do not have published karyotypes and four likely have missing chromosomes based on expectations from cytology, further highlighting the challenges of generating karyotype-complete genome assemblies for birds even when high-quality data is available and substantial manual curation time has been invested.

**Figure 2.**
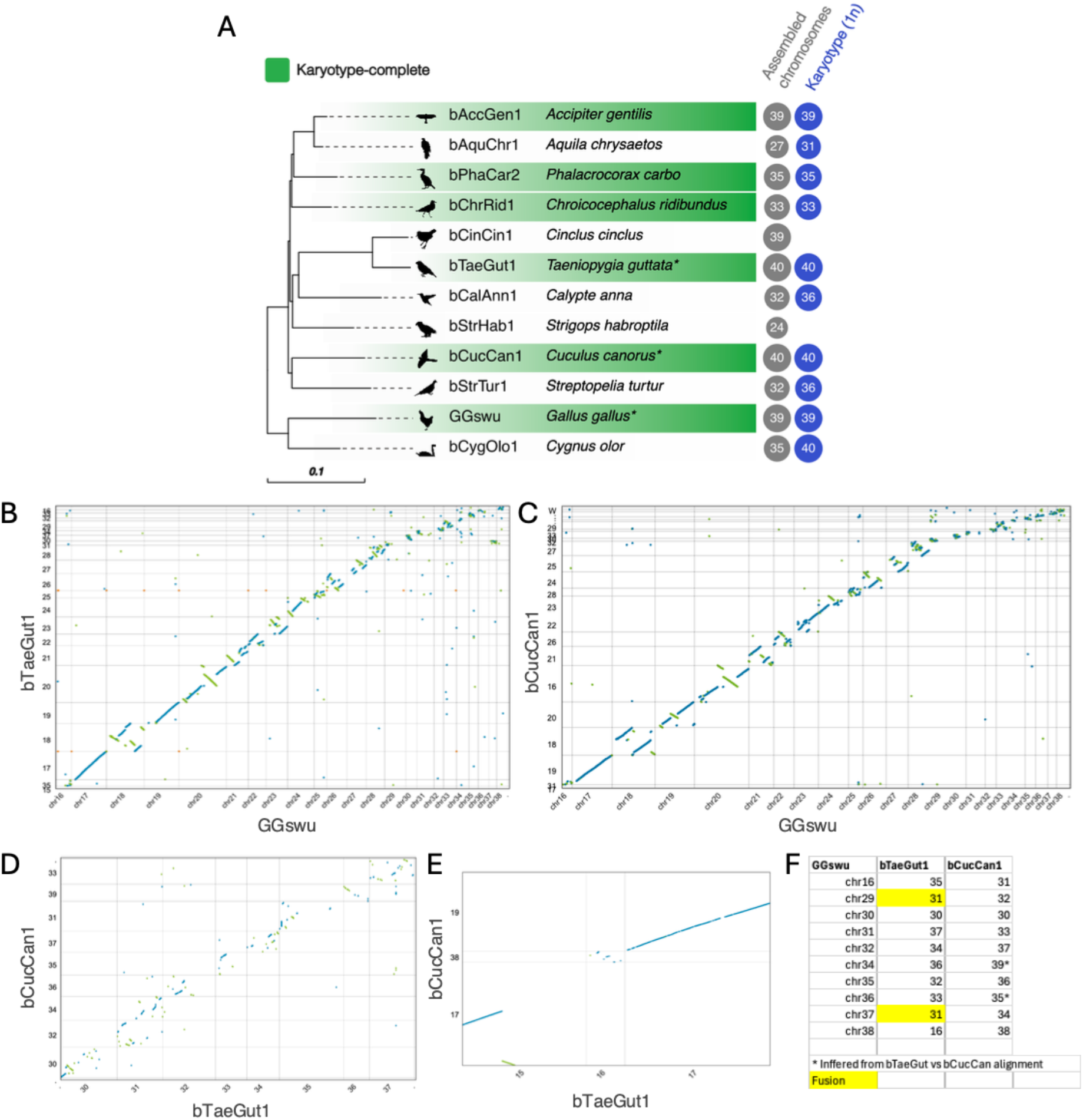
Phylogeny of annotated chromosome-scale bird reference genomes used to generate the MicroFinder protein set and conserved macro synteny of bird dot chromosomes. (**A**) Maximum likelihood phylogeny based on a concatenated alignment of 9,400 conserved single-copy orthogroups. Branch lengths are in amino acid substitutions per site. All nodes have ≥ 99% bootstrap support (1000 ultrafast bootstrap replicates). Species with genome assemblies confirmed to be karyotype-complete based on cytology are highlighted in green. Full details of all assemblies are given in **Supplementary Table 2.** PhyloPic (https://www.phylopic.org) silhouettes of each species are shown at the tree tips. Species marked with an “*” form the ToL reference set and are routinely used as references when assembling diverse bird genomes. (**B** -**F**) Dot chromosome synteny between genomes in the ToL reference set based on whole genome alignments. **F** summarises dot chromosome homology between GGswu, bTaeGut1 and bCucCan1 based on the alignments shown in **B** - **E**.

To identify conserved, low copy number genes to use as markers we clustered proteomes from the 12 bird reference genomes into orthogroups with OrthoFinder (Emms & Kelly, 2015, 2019) and used KinFin (Laetsch & Blaxter, 2017) to select broadly conserved “fuzzy” orthogroups that have relaxed conservation and copy number constraints (<= 3 gene copies per species and present in at least 50% of species). In total, 197,759 proteins were clustered into 16,589 orthogroups, of which 9,400 were conserved and single-copy in all species and 14,514 were identified by KinFin as “fuzzy” orthogroups (**Supplementary Table 3** and **Supplementary Data**). We further filtered the KinFin orthogroup set to only include genes located on dot chromosomes in any of the three ToL reference species, using the near-T2T GGswu chicken assembly to classify dot chromosome homologs in bCucCan1 and bTaeGut1 (**Figure 2B-F**). We reasoned that specifically targeting dot chromosomes rather than all microchromosomes would be most beneficial for assembly curation as larger microchromosomes are typically much less fragmented than dot chromosomes. This filtering identified 510 dot chromosome-associated orthogroups containing 4,510 proteins across all 12 reference species. To reduce redundancy, we clustered the dot chromosome-associated proteins with CD-HIT (Fu et al., 2012) to produce a final gene set containing 2,882 proteins which we refer to as the MicroFinder protein set.

Next, we investigated coverage of MicroFinder loci across, to our knowledge, the most complete bird genome assembled to date, the near-T2T GGswu assembly of chicken. The 10 GGswu dot chromosomes have between 15 and 67 GGswu MicroFinder loci per chromosome (307 in total), with an average density of 7.5 loci per Mb of sequence (**Figure 3**). In comparison, the orthoDB10 avian Benchmarking Universal Single-Copy Orthologs (BUSCO) gene set (n = 8,338 orthogroups) has only 3 genes located on dot chromosomes (**Supplementary Figure 1**), likely due historical difficulties with dot chromosome assembly leading to severe underrepresentation of dot chromosome genes in OrthoDB. Previously, Huang et. al. (2023) showed that chicken dot chromosomes are split into two distinct domains - gene-rich euchromatic regions and repetitive, gene-poor heterochromatic regions, with the euchromatic parts typically occupying a large region of the long arm of each chromosome. In line with this, we find clustering of MicroFinder proteins in high expression, low repeat density regions of dot chromosomes (**Figure 3**). As such, the high density of MicroFinder proteins in euchromatin will increase the likelihood of identifying genic regions of dot chromosomes in fragmented genome assemblies.

**Figure 3.**
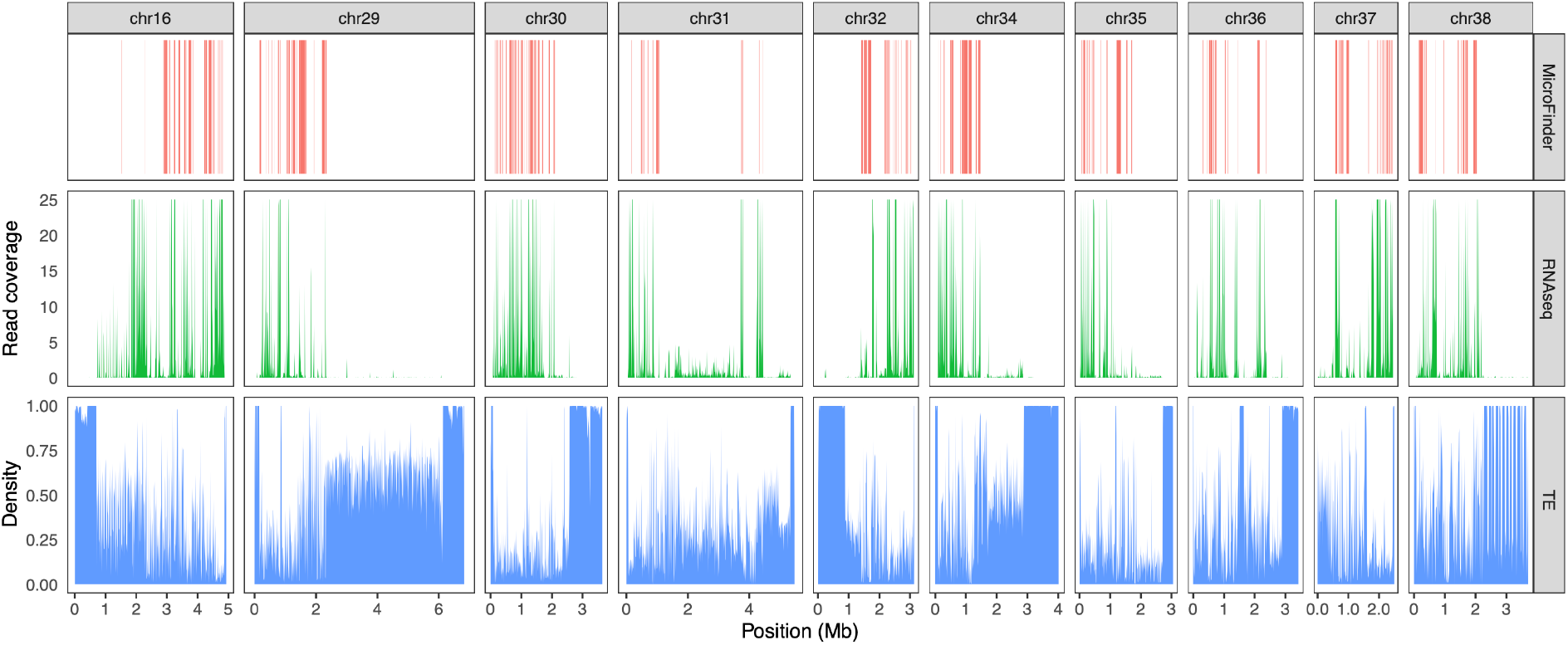
Distribution of MicroFinder proteins on chicken (GGswu assembly) dot chromosomes. Panels from top to bottom show the location of MicroFinder loci (coral), RNA-seq alignment counts from female chicken liver (SRR18788805) (green) in 10 Kb fixed windows, and transposable element density in 10 Kb fixed windows (blue). To aid visualisation of lower coverage genes, maximum RNAseq read coverage was capped at 25x.

### Gene mapping and assembly ordering to aid genome curation

To make use of the MicroFinder protein set we developed a pipeline to map and count MicroFinder proteins in a draft genome assembly and reorder scaffolds by MicroFinder protein count. This strategy means that putative dot chromosome scaffolds appear at the beginning of the Hi-C contact map separated from other small fragments, enabling curators to quickly identify dot chromosome content and start building up chromosome-scale scaffolds without having to sift through repetitive “shrapnel” contigs as is the case for a standard, size-sorted map. The MicroFinder pipeline aligns the MicroFinder protein set to a draft assembly with miniprot (H. Li, 2023), selects the top ranking hit for each protein, removes alignments with less than 70% identity and then counts protein alignments per scaffold and outputs a reordered assembly fasta file and associated MicroFinder count data. Optionally, the pipeline can apply a maximum scaffold size cutoff for assembly sorting. During testing we found that macrochromosome scaffolds can sometimes contain a low number of MicroFinder hits, most likely due to the presence of divergent paralogs or mis-mapping. We therefore recommend using a 5 Mb maximum scaffold size cutoff for assembly sorting. Following sorting, new Hi-C contact maps can be made for assembly curation in PretextView (https://github.com/sanger-tol/PretextView) using the CurationPretext pipeline (Pointon, 2025). MicroFinder has been packaged up into Docker and Singularity containers for easy deployment (https://github.com/sanger-tol/MicroFinder) and we have developed a training workshop with example datasets to guide users (Mathers et al., 2024).

To demonstrate how MicroFinder can be used as a curation aid, we applied it to the draft (pre curation) DToL genome assembly of *Anas acuta* (O’Brien & Lopez Colom, 2024). MicroFinder identified 61 putative dot chromosome scaffolds shorter than 5 Mb and moved them to the start of the Hi-C contact map (**Figure 4**). These scaffolds were manually ordered and rearranged to form 10 chromosomal elements during curation. Notably, we did not observe false positive MicroFinder ordered scaffolds with Hi-C signal placing them with macrochromosomes, indicating that MicroFinder proteins are reliable dot chromosome markers. This is likely due to conservation of microchromosome gene content and limited rearrangements between microchromosomes and macrochromosomes during avian evolution. As such, MicroFinder enables rapid curation of dot chromosomes using gene-rich scaffolds as anchors to build up dot chromosomes, removing the need for curators to trawl through repetitive shrapnel contigs and reducing the risk of small gene-rich dot chromosome contigs being missed from dot chromosome models during the curation process.

**Figure 4.**
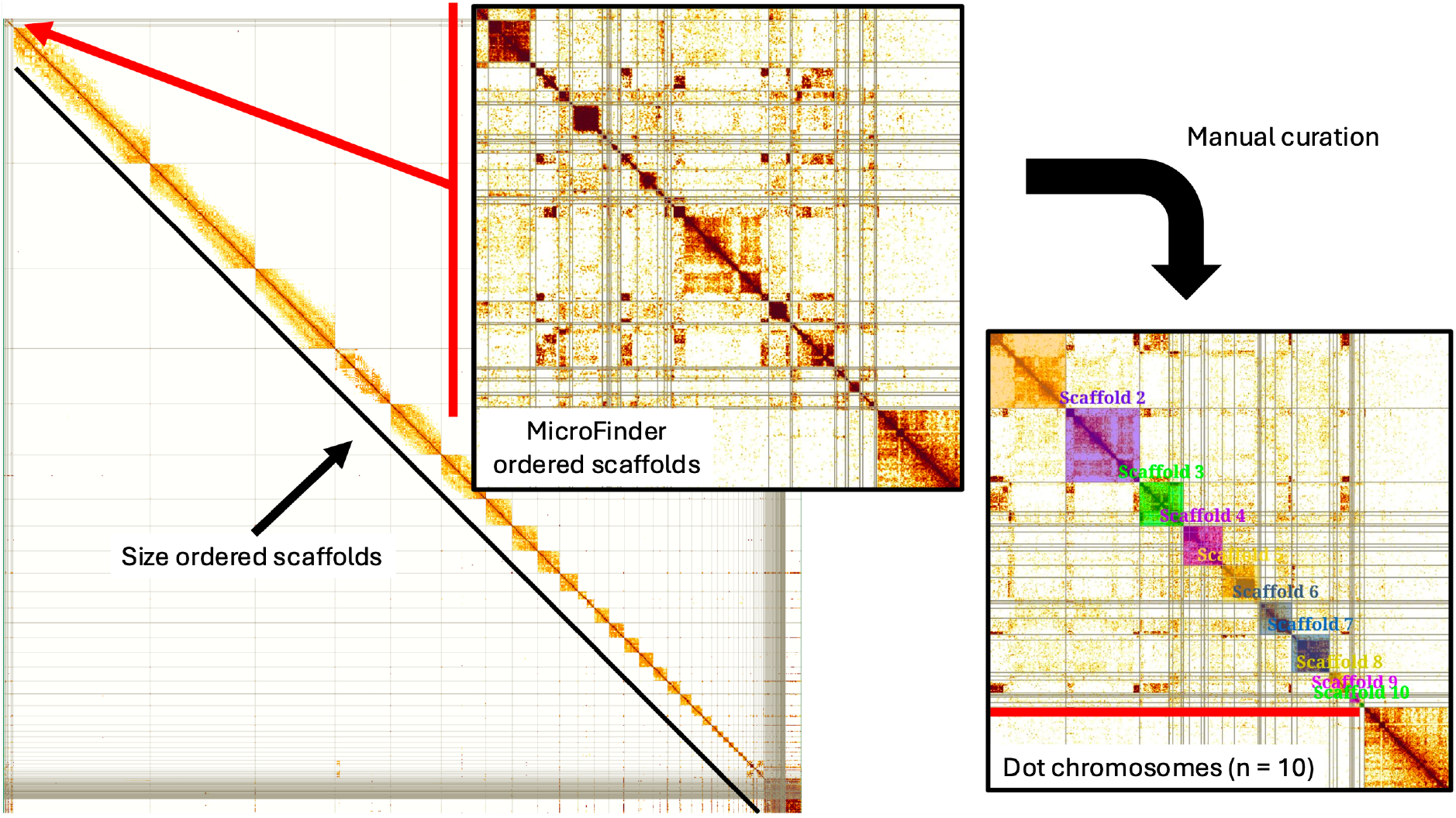
MicroFinder-enabled manual curation of bird dot chromosomes. Main panel shows Hi-C contact map of the MicroFinder-ordered draft (pre curation) genome assembly of *Anas acuta* (O’Brien & Lopez Colom, 2024). *Central panel* shows a zoomed in view of the putative dot chromosome content that has been moved to the start of the of the assembly by MicroFinder for curation. *Right hand panel* shows zoomed in view of the curated dot chromosomes.

### Reassembly of DToL bird genomes using MicroFinder-aided curation

Next, we investigated whether MicroFinder could be used to improve previously released chromosome-scale bird genome assemblies. We ran MicroFinder on 12 DToL bird genome assemblies that had been assembled using PacBio HiFi and Hi-C and subjected to manual curation by the ToL curation team (**Supplementary Table 4**). For each assembly, we ran MicroFinder with a 5 Mb maximum scaffold length cutoff and generated a new Hi-C contact map for curation in PretextView using the original sequence data. MicroFinder identified between 22 and 74 (mean = 49) putative unplaced dot chromosome scaffolds per assembly (**Figure 5a**). We were able to unambiguously place MicroFinder scaffolds onto dot chromosome models in 11 out of 12 of the assemblies, placing between 2 and 16 scaffolds and increasing the total length of assembled chromosomes in 9 out of 12 assemblies, adding between 216 Kb and 4.3 MB of additional content per assembly (average = 1.4 Mb) (**Figure 5b**). Two assemblies (bNetRuf1.1 and bAccGen1.1), had a decrease in assembled chromosome length due to identification of errors in the original assembly. In total, MicroFinder enabled the placement of an additional 12.5 MB of dot chromosome content across 9 DToL genomes. Furthermore, in the case of bAnaAcu1.1, were able to identify an additional dot chromosome model that had been missed in the original curation (**Figure 5c**). Unplaceable scaffolds either had ambiguous Hi-C signal or were too small to place, reflecting the fragmented nature of dot chromosome assemblies (**Figure 5c**).

**Figure 5.**
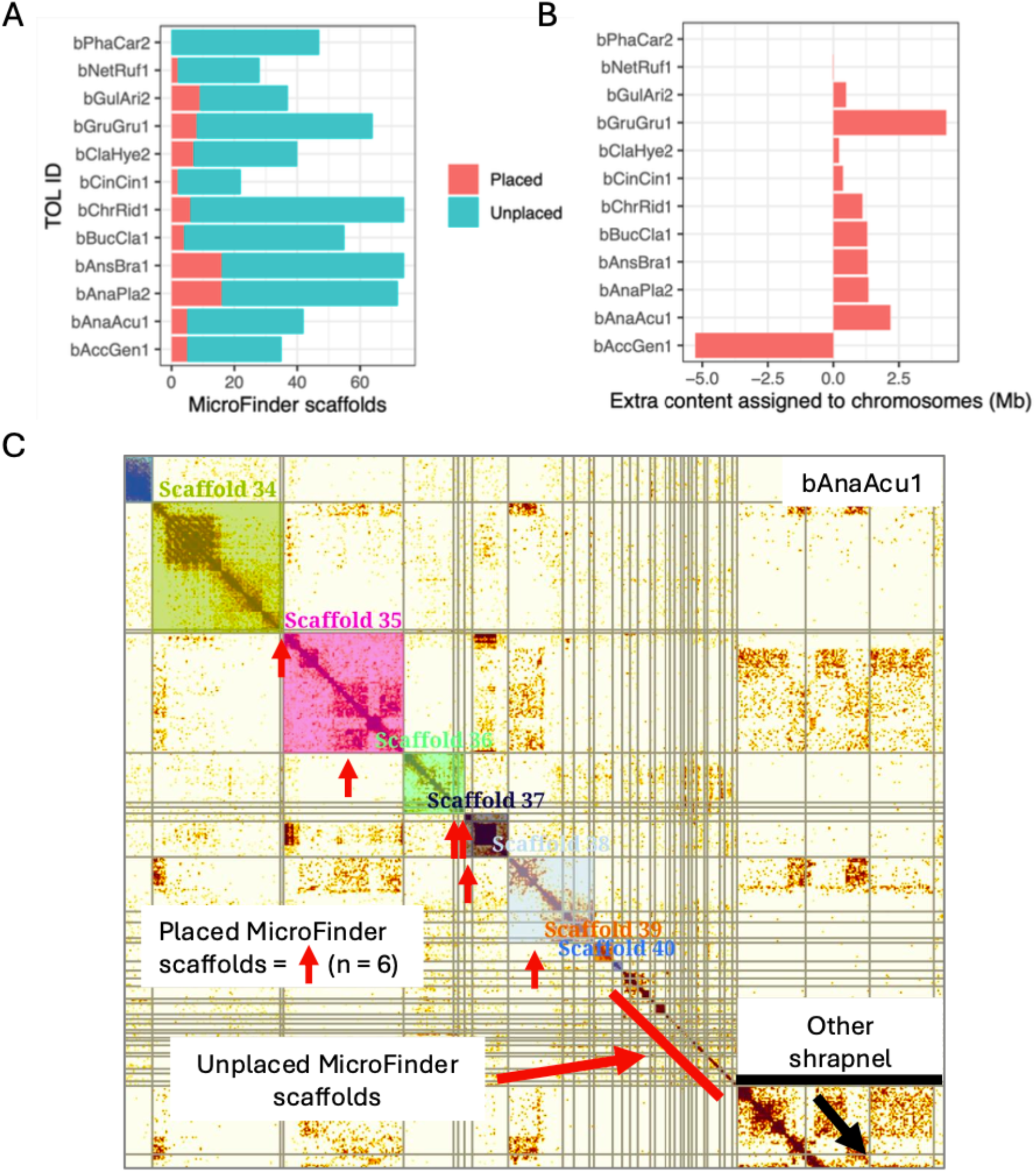
MicroFinder-enabled re-curation of 12 previously released DToL bird genome assemblies. (**A**) *Bar chart* showing counts of shrapnel scaffolds (previously unplaced content) identified by MicroFinder for 12 genome assemblies. Bars are coloured by whether the scaffolds were placed onto chromosome models during manual curation. (**B**) As for (**A**) but showing total sequence content added to chromosome models during manual curation of the MicroFinder sorted genome assemblies. (**C**) Hi-C contact map of the *Anas acuta* genome assembly (bAnaAcu1.1). The figure shows a zoomed in view of the smallest seven chromosomes. Scaffolds in the original assembly are separated by grey lines. Coloured squares indicate “painted” chromosomes and are assigned super scaffold IDs (Scaffold_(n)) by PretextView (shown above each square). Red vertical arrows indicate scaffolds that have been incorporated into chromosome models following MicroFinder-enabled manual re-curation. Scaffold_35 is a chromosome model that was unidentifiable in the original curation. Full stats for all 12 re-curated genome assemblies are provided in **Supplementary Table 4**.

## Conclusion

Here, we have identified a set of broadly conserved genes located on the smallest bird microchromosomes, known as dot chromosomes, and developed a pipeline (MicroFinder) to identify and order putative dot chromosome scaffolds in draft genome assemblies. By using “fuzzy” orthogroup selection, our gene set includes a large number of broadly conserved single-copy (or low copy number) genes and provides good coverage across all avian dot chromosomes (**Figure 3**). Using this strategy, MicroFinder can detect putative dot chromosome scaffolds in fragmented draft genome assemblies and is an effective curation aid for bird genome assembly, even enabling improvement to genome assemblies that have already undergone expert curation (**Figure 5**). Previously, an integrative method that uses a BAC panel to identify chromosome-specific regions was developed to resolve fragmented assemblies, including identification of microchromosomes (Damas et al., 2017), however it requires expertise in molecular cytogenetics and is time-consuming and impractical for current large-scale sequencing projects. Instead, MicroFinder provides a quick and easy pipeline to effectively pull-out putative dot chromosome fragments *in silico*. Furthermore, the MicroFinder approach may be applicable to other systems which have conserved but hard to assemble chromosomes, such as the dot chromosome (Muller element F) in Diptera.

Recently, near-T2T assemblies have been released for chicken, bustard and mallard (Hu et al., 2024; Huang et al., 2023; Luo et al., 2023). These assemblies achieved higher microchromosome contiguity through the inclusion of Oxford Nanopore ultra long reads. This approach represents a promising avenue to further improve bird genome assembly quality. However, due to scale and inertia, many projects still rely primarily on PacBio HiFi *de novo* assembly and will greatly benefit from our approach. We recommend MicroFinder is incorporated into bird genome assembly pipelines prior to manual curation to maximise the completeness of microchromosome assemblies.

## Methods

### Meta-analysis of bird karyotype and genome assembly chromosome counts

Genomes on a Tree (GoaT) (Challis et al., 2023) was used to retrieve bird chromosome counts based on cytology and from chromosome-level assemblies hosted INDC databases (**Supplementary Table 3**). Our query was made on the “taxon” index of the database, and we excluded taxa with missing data, retaining 105 species for downstream analysis. For chromosome counts based on genome assemblies, a single summary value was used as the representative chromosome count per species. For each assembly, the chromosome count corresponds to the number of chromosomes identified in the primary assembly (as opposed to the alternate assembly for a taxon). When multiple assemblies were available per taxon, the summary corresponds to the primary haplotype of NCBI RefSeq assembly. Haploid cytology-based chromosome numbers were extracted by halving the diploid number from the Bird Chromosome Database (Degrandi et al., 2020) and Animal Chromosome Counts Database (Release 1.0.1) (Román-Palacios et al., 2021) during GoaT import. A single summary value per species was calculated as the mode across all reported values per species. The ranges of values within each dataset were manually checked to ensure the summary values for chromosome number and haploid numbers from cytology were biologically consistent. We found that most of the variation detected within cytological observations corresponded to +-1 chromosome from the summary mode, consistent with reporting of different total number of chromosomes in different sexes and/or small miscounting from older manuscripts (e.g. Makino, 1951). The outliers were also manually checked on the original source, and all 7 detected cases corresponded to problematic values in their respective databases; because these values were not used as summaries, they were not included in our meta-analysis, and did not create bias in the data on **Figure 1a**. An interactive version of the scatterplot is available on the GoaT website for raw data exploration and download (https://tinyurl.com/4jnc3pbb).

### Dot chromosome homology assignment between GGswu, bTaeGut1 and bCucCan1

Pairwise whole genome alignments were carried out between chicken (GGswu), zebra finch (bTaeGut1) and cuckoo (bCucCan1) (**Supplementary Table 2**) using nucmer v4.0.0rc1 (Marçais et al., 2018) and visualised with Dot (https://dot.sandbox.bio/). Using these alignments, we identified homologs to GGswu dot chromosomes previously classified by Huang et al. (2023).

### Orthogroup clustering and identification of the MicroFinder protein set

To identify a set of conserved protein coding genes to use as dot chromosome markers we built orthogroups across representative bird genome assemblies. We selected 11 published chromosome-scale bird genome assemblies that had NCBI RefSeq or Ensembl rapid release gene-sets and combined them with a recent, near-T2T assembly of chicken (Huang et al., 2023) (**Supplementary Table 2**). For each species, we selected the longest transcript per gene to be the representative transcript and clustered protein sequences with OrthoFinder v2.5.4 (Emms & Kelly, 2015, 2019) in multiple sequence alignment mode (“-M msa”). The resulting orthogroups were filtered with KinFin v1.1.1 (Laetsch & Blaxter, 2017) with the parameters “--max 3 -x 0.5” to identify orthogroups present in at least 50 percent of species with a maximum of three gene copies per species. To create the MicroFinder protein set, the KinFin orthogroups were filtered to retain only those with a gene copy on chicken (GGswu), zebra finch (bTaeGut1) or cuckoo (bCucCan1) dot chromosomes. Proteins from the filtered orthogroups were then clustered with CD-HIT v4.8.1 (Fu et al., 2012) using default settings to reduce redundancy.

### Phylogenetic analysis

To place the 12 bird reference genomes used to generate the MicroFinder protein set in evolutionary context we carried out phylogenetic analysis using protein sequence alignments generated by OrthoFinder for 9,400 strictly conserved single-copy orthogroups. IQTree v2.3.4 was used to identify the optimal partitioning scheme, carry out model selection, estimate the maximum likelihood phylogeny and carry out 1,000 ultrafast bootstrap replicates to assess tree support (Chernomor et al., 2016; Kalyaanamoorthy et al., 2017; Minh et al., 2020a, 2020b, 2021). The IQTree phylogeny was rooted on the branch leading to Galloanserae (Galliformes plus Anseriformes) following Prum et al. (2015).

### The MicroFinder pipeline

All steps of the MicroFinder pipeline are implemented in a bash script and the whole pipeline is available as a docker or singularity container (https://github.com/sanger-tol/MicroFinder). First, the MicroFinder protein set is aligned to the draft genome assembly with miniprot v0.14 (H. Li, 2023) with default settings. From the resulting alignments, we retain the top hit and discard alignments with less than 70% identity. MicroFinder protein hits are counted for each scaffold and the input assembly fasta file is sorted by the alignment count. Optionally, a maximum scaffold length cutoff can be applied to the assembly sorting step. MicroFinder outputs a fasta file of the draft assembly sorted by MicroFinder protein alignment counts, a table of alignment counts per input scaffold and a GFF file of the miniprot alignments. It should be noted that MicroFinder counts reflect the number of protein hits from the MicroFinder protein set rather than counts of individual loci. We opted to map all proteins to maximise sensitivity to detect candidate dot chromosome scaffolds across a wide range of bird species. The MicroFinder-sorted assembly file should be prepared for manual curation in PretextView (https://github.com/sanger-tol/PretextView) with the CurationPretext pipeline (Pointon, 2025) with the “--no-sort” parameter used to retain the order of the MicroFinder assembly file in the Hi-C contact map.

### MicroFinder protein distribution in chicken (GGswu) and associated features

We investigated the distribution of MicroFinder proteins across the near-T2T GGswu chicken assembly (Huang et al., 2023). MicroFinder protein coordinates were extracted from the GGswu annotation GFF file. To place MicroFinder proteins in context we also estimated genome-wide repeat content and gene expression levels. RNA-seq from a from female chicken liver (SRR18788805) was aligned to the GGswu assembly with HISAT2 v2.2.1 (Kim et al., 2015) and we calculated read depth in 10 Kb fixed windows using Sambamba v0.8.2 (Tarasov et al., 2015). To estimate repeat density across the GGswu dot chromosomes, we ran RepeatMasker v4.1.8 (Smit et al., 2005; Tarailo-Graovac & Chen, 2009) using a manually curated avian repeat library (Peona et al., 2023; Peona, Palacios-Gimenez, et al., 2021; Pointon, 2025) and calculated repeat density in 10 Kb fixed windows with bedtools coverage v2.31.1 (Quinlan c49 Hall, 2010) using the RepeatMasker GFF file as input. To compare the distribution of MIcroFinder proteins to BUSCO genes we ran BUSCO v5.8.2 (Simão et al., 2015; Waterhouse et al., 2018) with the Aves OrthoDB gene set (n = 8338) on the GGswu assembly and extracted the coordinates of BUSCOs located on the dot chromosomes.

### Reassembly of DToL bird genomes with MicroFinder-enabled curation

We selected 12 previously published DToL bird genome assemblies for re-curation with MicroFinder (**Supplementary Table 4**). For each assembly, we ran MicroFinder with a 5 Mb maximum scaffold length cutoff and generated a new Hi-C contact map for curation in PretextView using the CurationPretext pipeline v1.0.1 (Pointon, 2025) with the “--no-sort” parameter. CurationPretext was provided with the original Hi-C and PacBio long reads for each assembly to create a Hi-C contact map with read coverage, gap, telomere and simple repeat density tracks. Manual curation was carried out using PretextView v1.0.0 (https://github.com/sanger-tol/PretextView). Following manual curation of each assembly, an AGP file was exported from PretextView and an updated assembly generated using pretext-to-asm (https://github.com/sanger-tol/agp-tpf-utils).

## Data and Code availability

Supplementary data containing OrthoFinder results, the MicroFinder gene set and the 12 re-curated bird genome assemblies is available from Zenodo (https://doi.org/10.5281/zenodo.15364993). For each of the re-curated genome assemblies, we have provided a MicroFinder-ordered Hi-C contact map of the original assembly, PretextView savestate and agp files to show changes made to the original assembly and an updated FASTA file of the assembly. The MicroFinder source code and containerised versions of the pipeline are available on GitHub (https://github.com/sanger-tol/MicroFinder). Mathers et. al. (2024) provides a practical guide for using MicroFinder-ordered assemblies for curation with example datasets (https://doi.org/10.5281/zenodo.13913870).

## Acknowledgments

We thank Prof. Alex Suh and Dr Valentina Peona for providing access to their curated avian repeat library. We thank Dr Kerstin Howe and Kr Kamil Joran for comments on an earlier version of the manuscript. This work was supported by Wellcome through core funding to the Wellcome Sanger Institute (220540) and the Darwin Tree of Life Discretionary Award (218328).

