## Supplementary Figure 1 for "MicroFinder: Conserved gene-set mapping and assembly ordering for manual curation of bird microchromosomes"

### Supplementary Figures

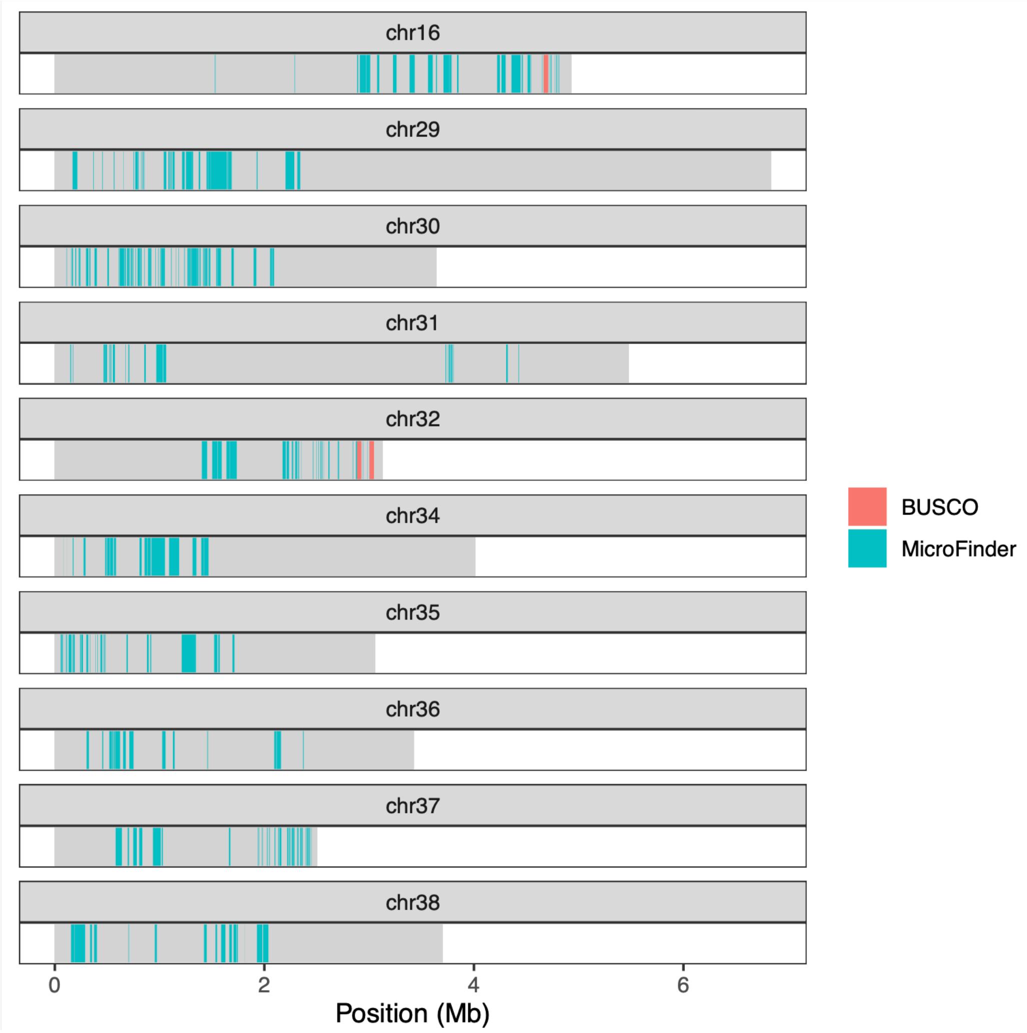


**Supplementary Figure 1:** Location of orthoDB10 avian BUSCOs (n = 3) and MicroFinder loci (n = 307) on chicken (GGswu assembly) dot chromosomes.
